# RainBar: Optical Barcoding for Pooled Live-Cell Imaging with Single-Cell Resolution

**DOI:** 10.1101/2025.11.04.686676

**Authors:** Ruzbeh Mosadeghi, Daniel Foyt, Louis Sharp, Catherine Taylor, Neil Tay, Stefan Oberlin, John Fan, Struan Bourke, Michael Kattah, Bo Huang, Michael T McManus

## Abstract

High-throughput pooled screening has advanced functional genomics, but most existing methods rely on endpoint sequencing and are blind to dynamic, time-resolved phenotypes. We developed RainBar (Rainbow Barcodes), an optical barcoding system that supports pooled live-cell imaging with single-cell resolution. RainBar uses lentiviral co-delivery of spectrally distinct nuclear and cytoplasmic fluorescent proteins to encode up to 64 unique perturbations per well. To mitigate barcode recombination and improve decoding accuracy, we employed single-template viral production, codon diversification, and a ratio-based spectral unmixing pipeline tailored to overlapping fluorophores. An inverted cytoplasmic segmentation approach and multilayer perceptron classifier enabled accurate barcode identification in both arrayed and pooled formats. As a proof of concept, we applied RainBar to dissect NF-κB signaling dynamics in epithelial cells. Live imaging of RelA translocation uncovered stimulus-specific kinetics: IL-1β triggered rapid recovery, while TNF induced prolonged nuclear localization. In pooled CRISPRi screens, RainBar recovered known NF-κB regulators (e.g., IL1R1, MYD88, TNFRSF1A) and highlighted additional modulators, including the Ino80 chromatin remodeling complex subunits and KAT2A acetyltransferase. Together, these results position RainBar as a flexible platform for multiplexed, image-based functional genomics, with potential to reveal dynamic signaling architectures across diverse cellular contexts in live cells.

## INTRODUCTION

Pooled CRISPR screening has transformed functional genomics by enabling systematic dissection of gene function in mammalian cells^1–3^. Most pooled screens, however, rely on sequencing-based readouts, an approach that, while scalable and mature, is fundamentally destructive. These assays readily capture endpoint phenotypes such as viability, gene expression, or reporter abundance, but fall short when it comes to more complex or dynamic phenotypes, such as changes in morphology, signaling dynamics, or subcellular localization. Optical pooled screening (OPS) has emerged to bridge this gap, allowing imaging-based phenotypes to be linked to perturbations using barcodes readable via in situ hybridization, in situ sequencing, or antibody^4–6^. These approaches have shown compelling results in fixed cells and in some cases, intact tissues, but they are generally incompatible with live imaging, limiting their utility for studying dynamic or reversible processes.

Stochastic multicolor labeling strategies like Brainbow can generate hundreds of unique barcodes by random recombination, but this randomness makes it difficult to assign a unique perturbation to a specific barcode without large-scale oversampling^7,8^. This becomes especially problematic in live-cell imaging experiments that require repeated sampling over time. In these settings, achieving sufficient barcode resolution demands high cell numbers and uniform barcode representation; neither of which are trivial when using compact viral vectors like lentivirus or AAV^9,10^ Combinatorial approaches that encode information via localization patterns or tag length have expanded the barcode design space, but suffer from inconsistent marker performance across cell types, or skewed barcode distributions due to inefficient packaging or expression. For example, visually encoded systems like EpicTags and visual barcodes rely on localization signals that may not function uniformly across lines, and typically offer limited temporal resolution^6,11^. Even deterministic barcoding systems, (such as those based on photoactivatable proteins) remain constrained to single phenotypes or lack temporal depth^12–17^. Taken together, creating a barcode system that is simultaneously deterministic, scalable, and compatible with live imaging remains a substantial challenge, complicated by practical constraints. For example, the limited payload size for AAV restricts barcode sizes^9^. Recombination-prone lentiviral delivery can decouple guides from barcodes^10,18^. Meanwhile, standard segmentation strategies utilizing separate markers for each subcellular compartments competes for the limited number of color channels.

Here we present RainBar (Rainbow Barcodes), a deterministic optical barcoding platform that encodes perturbations via predefined combinations of spectrally distinct fluorescent proteins (FPs) targeted to the nucleus or cytoplasm. Each barcode is intentionally designed, rather than stochastically generated, enabling 64 uniquely identifiable perturbations per well without requiring oversampling. RainBar uses lentiviral co-delivery of nuclear and cytoplasmic FPs, with codon-diversified sequences and a single-template production workflow to minimize recombination, an inverted cytoplasmic masking strategy for segmenting two compartments with a single marker, and a ratio-based unmixing pipeline to resolve spectrally overlapping fluorescent proteins^19,20^.

As a proof of concept, we applied RainBar to monitor NF-κB signaling dynamics in epithelial cells, focusing on stimulus-specific patterns of RelA translocation. In pooled CRISPRi screens, RainBar recovered canonical NF-κB pathway regulators (including IL1R1, MYD88, and TNFRSF1A) and highlighted additional modulators such as the Ino80 chromatin remodeling complex subunits and the histone acetyltransferase KAT2A. These results position RainBar as a generalizable platform for multiplexed, image-based pooled screening, particularly well-suited for dissecting dynamic signaling events in live cells.

## RESULTS

### The design of RainBar barcodes

We set out to develop a reproducible, multiplexed FP barcoding system (RainBar) that would support live-cell imaging at short acquisition intervals while scaling to dozens of perturbations per well. Each RainBar construct encodes an sgRNA under a human U6 promoter and a two-part FP barcode under the control of EF1α: one fluorophore directed to the nucleus (via an NLS) and the other to the cytoplasm (via an NES), separated by a P2A peptide linker (Fig. 1A,B). Barcodes are denoted in the format NLS-FP1– NES-FP2 (e.g., EGFP–mCherry indicates nuclear EGFP and cytoplasmic mCherry).

**Figure 1.**
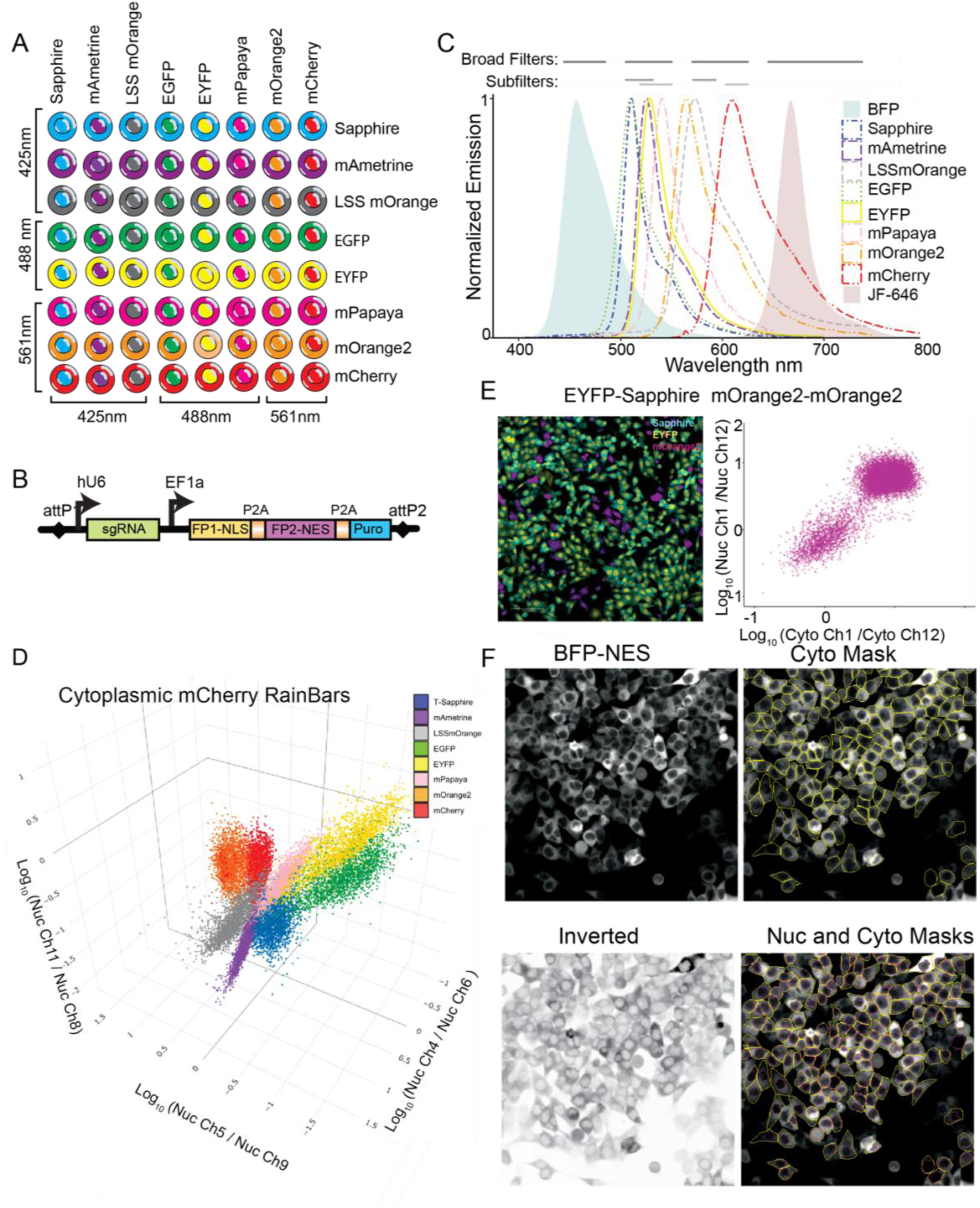
Design of the RainBar barcode system, segmentation strategy, and minimization of lentiviral recombination. **a,** The RainBar-64 library consists of eight spectrally distinct fluorescent proteins (FPs) targeted to the nucleus (NLS) and cytoplasm (NES), yielding 64 deterministic barcode combinations. Numbers indicate excitation wavelengths used for each FP. **b,** Lentiviral transfer vector architecture showing an sgRNA under a human U6 promoter and a bicistronic barcode cassette under EF1α control, with nuclear and cytoplasmic FPs separated by a P2A sequence. A puromycin resistance gene enables selection. **c,** Emission spectra of all RainBar FPs (dashed lines), including the segmentation marker BFP and assay fluorophore JF-646 (shaded), alongside filter configurations used on the Opera Phenix. Wide filters (top) and subfilters (bottom) were used for ratio-based spectral unmixing. **d,** Ratio-based unmixing of nuclear intensity signals enables robust separation of all eight NLS-tagged FPs when paired with cytoplasmic mCherry. Shown are three representative log-scale nuclear intensity ratios. **e,** Pooled virus produced using PRECISE lentiviral packaging with codon-diversified constructs minimizes barcode recombination. Representative 2D ratio feature plots show distinct clustering for EYFP–Sapphire and mOrange2–mOrange2 barcodes. **f,** Inverted segmentation approach for simultaneous nuclear and cytoplasmic masking. BFP-NES fluorescence enables cytoplasmic segmentation with Cellpose (top). The same images, inverted in contrast, allow nuclear segmentation from the same marker (bottom). Combined cytoplasmic (yellow) and nucleus (magenta).

While larger barcode spaces can be constructed using additional FPs or subcellular localizations, we found that real-time live imaging places practical constraints on barcode complexity. Rather than attempting tens of thousands of barcodes in a single well, we focused on smaller sub-pools that remain tractable for high-frame-rate acquisition. Using eight distinct FPs across two compartments, we designed a 64-barcode set, which is sufficient to support pooled imaging of ∼20,000–40,000 cells per 96-well format, yielding ∼300–600 cells per perturbation. This density allows for kinetic analyses at single-cell resolution. In non-live-imaging contexts, alternative localizations (e.g., plasma membrane, mitochondria, peroxisomes) and expanded FP palettes could scale this approach to barcode libraries exceeding 20,000 combinations^21–23^. Genome-scale screens are feasible with the RainBar-64 set across 4 to 8 96-well plates, or 1 to 2 384-well plates, depending on coverage requirements and replicate design. One limitation, however, is that RainBar barcodes are not ideally suited to assays that directly perturb nuclear import/export pathways, as such perturbations may confound localization signals. In these cases, cytoplasmic FPs paired with non-nuclear localizations are preferred.

A core challenge in constructing this barcode set was reliable spectral separation of overlapping fluorophores. We dedicated the BFP channel for segmentation (via nuclear exclusion), and farred fluorophores (e.g., JF-646^24^) for biological readouts, leaving the green–yellow–red window for the barcodes (Fig. 1C). Within this spectral window, robust multiplexing was achieved by four strategies. First, we additionally utilized fluorescent proteins with large Stokes shifts, allowing for divergent emission despite shared excitation sources^21–23,25^. For example, both T-Sapphire (Sapphire) and LSSmOrange are efficiently excited by the 425 nm laser used for TagBFP (BFP), but exhibit distinct emission profiles (Fig. S1A)^21–23,25^. Second, FP signal varied considerably due to both intrinsic brightness differences and excitation wavelength match (e.g., mPapaya is intrinsically bright but is only ∼39% excited by the 488 nm laser). Therefore, for some channels, we captured both high- and low-exposure images to expand the dynamic range (Table 1). Third, we used a combination of wideband emission filters and narrow band subfilters on the Opera Phenix high content microscope to discriminate signals from FPs with high spectral overlap (e.g. mOrange2 and mCherry) (Fig. 1C). Lastly, to account for the variability of absolute expression levels, we extracted relative ratios of fluorescence signal across multiple filters, with remain stable across a wide range of expression level (Fig. S1B,E) ^19^. Using this approach, we demonstrated the successful discrimination of all eight FPs in the nuclear compartment when paired with a constant cytoplasmic FP (Fig. 1D;).

**Table 1.** Excitation and emission settings used for RainBar barcode decoding on the Opera Phenix platform. Excitation lasers, emission filters, exposure times, and excitation power percentages are listed for each channel used in ratio-based spectral unmixing. Channels were selected to minimize spectral bleed-through while maximizing signal-to-noise.

| CHANNEL | EXCITATION LASER (NM) | EMISSION FILTER (NM) | POWER | EXPOSURE TIME (MS) | EXCITATION POWER (%) |
| --- | --- | --- | --- | --- | --- |
| 1 | 425 | 435-480 | Low | 100 | 60 |
| 2 | 425 | 500-550 | Low | 400 | 100 |
| 3 | 425 | 500-530 | Low | 400 | 100 |
| 4 | 425 | 515-550 | Low | 400 | 100 |
| 5 | 425 | 570-630 | Low | 400 | 100 |
| 6 | 488 | 500-550 | Low | 100 | 50 |
| 7 | 488 | 500-550 | High | 400 | 100 |
| 8 | 488 | 500-530 | Low | 100 | 50 |
| 9 | 488 | 500-530 | High | 400 | 100 |
| 10 | 488 | 515-550 | Low | 100 | 50 |
| 11 | 488 | 515-550 | High | 400 | 100 |
| 12 | 561 | 570-630 | Low | 200 | 100 |
| 13 | 561 | 571-596 | Low | 200 | 100 |
| 14 | 561 | 605-630 | Low | 200 | 100 |

### Minimizing lentiviral recombination for homologous barcodes

In pooled CRISPR screens, lentiviral delivery of sgRNAs and barcodes from plasmid pools introduces a persistent challenge: recombination during reverse transcription can decouple guides from their intended barcodes^10^. This issue is often masked in sequencing-based screens, where barcode–guide associations can be computationally recovered. However, in optical systems like RainBar (where fluorescent proteins are the sole readout) such mismatches are irreversible, and even rare recombination events can confound interpretation.

To reduce these events, we implemented a single-template lentiviral production strategy based on the PRECISE system (Tay et al., manuscript in preparation). In this system, HEK293T packaging cells stably express lentiviral elements and carry a recombinase-addressable landing pad at the AAVS1 locus (Fig. S1C). Library plasmids encoding both the sgRNA and the FP barcode are flanked by attP sites and stably integrated into the AAVS1 site, such that each producer cell harbors only one transfer vector. This dramatically reduces the likelihood of barcode–guide decoupling, since co-packaging of multiple RNA genomes (an upstream driver of recombination) is effectively eliminated. In this configuration, any residual recombination is more likely to yield non-functional or silent variants than to produce mismatched barcodes.

Despite this improvement, for many RainBar constructs we only observed a FP signal in the cytoplasm. Interestingly, it was the FP assigned to the nucleus that was expressing in the cytoplasm, while no FP signal was noted in the nucleus. This was particularly a problem with highly homologous FP pairs (EGFP–EGFP >EGFP–EYFP > EGFP–mCherry) (Fig. S1D,G). These asymmetric losses suggested a homology-driven recombination mechanism affecting NLS-FP coding sequence^10^. To mitigate this issue, we codon-diversified each fluorescent protein to reduce nucleotide-level sequence identity between the nuclear and cytoplasmic FP variants while preserving their amino acid sequence (Fig. S1F). Because residues like tryptophan and methionine are constrained to single codons, we did not attempt complete elimination of homology but, instead, made broad reductions to keep shared stretches under 7 nucleotides (nt), with only a few cases up to 14 nt. We quantified nuclear FP retention using the nuclear-to-cytoplasmic (N/C) intensity ratio of the relevant channel. Codon diversification significantly reduced nuclear signal loss (Fig. S1D,E,G), although very rare mislocalization events persisted for all RainBars (eg. such as cytoplasmic-only EYFP) (Fig. 1E)^26^.

Codon-diversification slightly affected signal strength. DsRed-derived proteins remained similarly robust after optimization, but the first EGFP variant (EGFP-A) showed reduced fluorescence. A second design (EGFP-B), generated using the Twist codon optimization algorithm, restored sufficient brightness for imaging (Fig. S2B). To balance performance across the barcode library, we placed the codon-diversified FP in the NLS position and retained the canonical version in the NES position (Fig. 1B), optimizing for both recombination resistance and fluorescence intensity.

### Simultaneous nuclear and cytoplasmic segmentation using a single marker

Reliable segmentation is a prerequisite for pooled live-cell imaging, but standard nuclear markers present unexpected tradeoffs in this context. Dyes like Hoechst stain nuclei efficiently and are widely used for segmentation, yet their spectral properties pose challenges here: Hoechst emits brightly and broadly in the violet range, overlapping significantly with barcoding fluorophores such as Sapphire and complicating spectral separation (Fig. S2B)^27^. We explored alternatives, including SPY700-DNA, but found that staining quality varied between cells, limiting its utility as a consistent segmentation marker (Fig. S2B)^28^. Moreover, DNA dyes by design only label nuclei, offering no information about cytoplasmic boundaries.

To sidestep these issues, we constructed a BFP-NES fusion expressed under EF1α to serve as a uniform cytoplasmic marker. Cytoplasmic masks were generated using Cellpose, which segments the entire cell based on the BFP signal. We then inverted the raw image data, transforming the dark nuclei into bright islands. Using standard nuclear segmentation parameters, Cellpose readily detects these nuclei while rejecting similarly dark regions between cells based on the shape (Fig. 1F)^20^.

We compared two versions of the BFP-NES construct: one linked to a Bleomycin resistance gene and the other to Hygromycin. Both enabled cytoplasmic and nuclear segmentation, but the Hygromycin-linked version exhibited dimmer BFP fluorescence. This lower intensity reduced spectral interference with green and yellow barcoding channels, while still providing sufficient signal for reliable segmentation (Fig. S2D). That said, violet excitation (used to excite BFP) is known to be phototoxic in some cell types– especially during long-term time-lapse imaging^27^. In such cases, the brighter BFP-NES-Bleo construct may be preferable, particularly when dealing with weakly transduced cells or assays requiring extended imaging.

### Decoding Rainbar

To establish a reference dataset for barcode classification, we individually plated all 64 RainBar constructs across a 96-well plate and acquired images under 14 laser/filter configurations (11 distinct combinations plus 3 repeated at higher exposure (Table 1)). After segmentation, we applied quality filters to exclude very small or overly circular masks, which typically reflect dying cells or debris. Only nuclei fully enclosed within a cytoplasmic mask were retained for downstream analysis. We trimmed the masks to minimize contaminations between the nuclear signal, cytoplasmic signal and signal from neighboring cells (see Methods).

To decode the full 64-barcode library, we calculated all pairwise fluorescence ratios among the 14 imaging channels for each subcellular compartment independently. These features were then used to train a multilayer perceptron (MLP) classifier, using 70% of cells from single-construct wells for training and 30% for testing (Fig. 2A)^29^. The MLP architecture consisted of four hidden layers, each with 64 neurons, and was trained for 10 epochs. A supervised UMAP projection of the test set revealed clean separation of the full barcode space, indicating strong feature discriminability (Fig. 2B).

**Figure 2.**
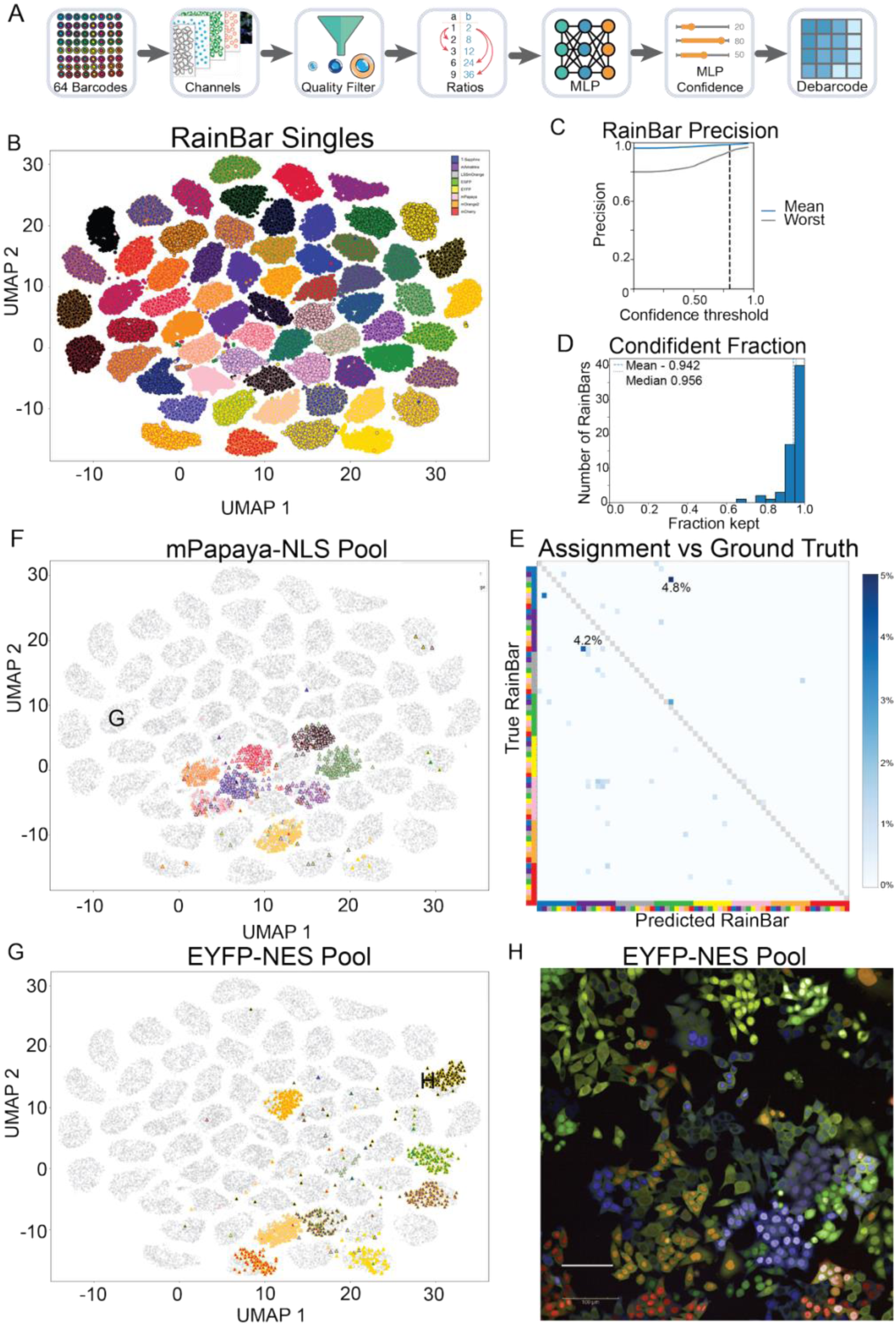
Barcode classification performance and pooled decoding using RainBar. **a,** Schematic of the RainBar decoding pipeline. Cells from single-barcode wells were imaged across 14 filter settings and segmented; pairwise intensity ratios were extracted separately for nuclear and cytoplasmic channels, then used to train a multilayer perceptron (MLP) classifier. **b,** Supervised UMAP embedding of the held-out test set (30% of single-barcode cells) shows clear separation of all 64 barcodes. Fill color indicates NLS FP identity; border color indicates NES FP identity. **c,** Per-barcode classification precision, with mean across barcodes in blue and the lowest-performing barcode (Sapphire–LSSmOrange) in gray. Dashed line indicates confidence threshold of 0.8. **d,** Distribution of the fraction of cells retained after filtering by classification confidence ≥0.8 across all barcodes (mean = 94.2%, median = 95.6%). **e,** Confusion matrix comparing MLP-assigned versus true barcodes, ordered by NLS (left/bottom); NES (right/top) based on legend in panel B (Note: color scale capped at 5% to accentuate rare errors. Identity as in b. Color scale capped at 5% to emphasize rare misclassifications; most errors were low-frequency and asymmetrical. **f,** Pooled decoding: cells from a validation pool containing all eight mPapaya-NLS RainBars are overlaid on the reference UMAP from b. **g,** Same as f, for a pool containing all eight EYFP-NES barcodes. **h,** Representative pooled image from the EYFP-NES validation.

Classification performance was high. Without filtering on MLP prediction confidence, the average precision across barcodes was 96.4%. Applying a confidence threshold of ≥ 0.8 retained over 95.6% of quality-filtered cells, with mean precision, recall, and F1 scores of 98.7%, 92.8%, and 95.6%, respectively (Fig. 2C,D; Table 2). With this filtering, even the lowest performing barcode (Sapphire–LSSmOrange) achieved 96.1% precision and a recall of 63.5%, corresponding to an F1 score of 74.6%.

**Table 2.**
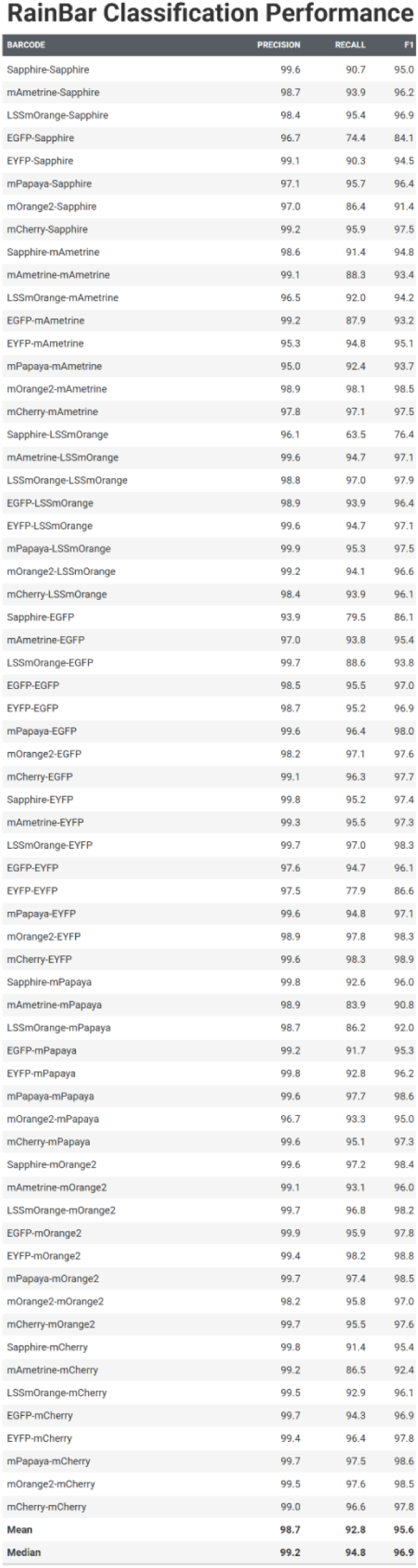
Classification accuracy for RainBar single-construct barcodes. Precision, recall, and F1 scores were computed for each of the 64 barcode combinations in the RainBar library. Scores are based on a multilayer perceptron (MLP) classifier trained on held-out single-barcode wells. Each barcode comprises a unique pair of NLS- and NES-targeted fluorescent proteins. Barcodes are listed in NLS–NES format. Mean and median values across the 64 barcodes are shown at bottom.

| BARCODE | PRECISION | RECALL | F1 |
| --- | --- | --- | --- |
| Sapphire-Sapphire | 99.6 | 90.7 | 95.0 |
| mAmetrine-Sapphire | 98.7 | 93.9 | 96.2 |
| LSSmOrange-Sapphire | 98.4 | 95.4 | 96.9 |
| EGFP-Sapphire | 96.7 | 74.4 | 84.1 |
| EYFP-Sapphire | 99.1 | 90.3 | 94.5 |
| mPapaya-Sapphire | 97.1 | 95.7 | 96.4 |
| mOrange2-Sapphire | 97.0 | 86.4 | 91.4 |
| mCherry-Sapphire | 99.2 | 95.9 | 97.5 |
| Sapphire-mAmetrine | 98.6 | 91.4 | 94.8 |
| mAmetrine-mAmetrine | 99.1 | 88.3 | 93.4 |
| LSSmOrange-mAmetrine | 96.5 | 92.0 | 94.2 |
| EGFP-mAmetrine | 99.2 | 87.9 | 93.2 |
| EYFP-mAmetrine | 95.3 | 94.8 | 95.1 |
| mPapaya-mAmetrine | 95.0 | 92.4 | 93.7 |
| mOrange2-mAmetrine | 98.9 | 98.1 | 98.5 |
| mCherry-mAmetrine | 97.8 | 97.1 | 97.5 |
| Sapphire-LSSmOrange | 96.1 | 63.5 | 76.4 |
| mAmetrine-LSSmOrange | 99.6 | 94.7 | 97.1 |
| LSSmOrange-LSSmOrange | 98.8 | 97.0 | 97.9 |
| EGFP-LSSmOrange | 98.9 | 93.9 | 96.4 |
| EYFP-LSSmOrange | 99.6 | 94.7 | 97.1 |
| mPapaya-LSSmOrange | 99.9 | 95.3 | 97.5 |
| mOrange2-LSSmOrange | 99.2 | 94.1 | 96.6 |
| mCherry-LSSmOrange | 98.4 | 93.9 | 96.1 |
| Sapphire-EGFP | 93.9 | 79.5 | 86.1 |
| mAmetrine-EGFP | 97.0 | 93.8 | 95.4 |
| LSSmOrange-EGFP | 99.7 | 88.6 | 93.8 |
| EGFP-EGFP | 98.5 | 95.5 | 97.0 |
| EYFP-EGFP | 98.7 | 95.2 | 96.9 |
| mPapaya-EGFP | 99.6 | 96.4 | 98.0 |
| mOrange2-EGFP | 98.2 | 97.1 | 97.6 |
| mCherry-EGFP | 99.1 | 96.3 | 97.7 |
| Sapphire-EYFP | 99.8 | 95.2 | 97.4 |
| mAmetrine-EYFP | 99.3 | 95.5 | 97.3 |
| LSSmOrange-EYFP | 99.7 | 97.0 | 98.3 |
| EGFP-EYFP | 97.6 | 94.7 | 96.1 |
| EYFP-EYFP | 97.5 | 77.9 | 86.6 |
| mPapaya-EYFP | 99.6 | 94.8 | 97.1 |
| mOrange2-EYFP | 98.9 | 97.8 | 98.3 |
| mCherry-EYFP | 99.6 | 98.3 | 98.9 |
| Sapphire-mPapaya | 99.8 | 92.6 | 96.0 |
| mAmetrine-mPapaya | 98.9 | 83.9 | 90.8 |
| LSSmOrange-mPapaya | 98.7 | 86.2 | 92.0 |
| EGFP-mPapaya | 99.2 | 91.7 | 95.3 |
| EYFP-mPapaya | 99.8 | 92.8 | 96.2 |
| mPapaya-mPapaya | 99.6 | 97.7 | 98.6 |
| mOrange2-mPapaya | 96.7 | 93.3 | 95.0 |
| mCherry-mPapaya | 99.6 | 95.1 | 97.3 |
| Sapphire-mOrange2 | 99.6 | 97.2 | 98.4 |
| mAmetrine-mOrange2 | 99.1 | 93.1 | 96.0 |
| LSSmOrange-mOrange2 | 99.7 | 96.8 | 98.2 |
| EGFP-mOrange2 | 99.9 | 95.9 | 97.8 |
| EYFP-mOrange2 | 99.4 | 98.2 | 98.8 |
| mPapaya-mOrange2 | 99.7 | 97.4 | 98.5 |
| mOrange2-mOrange2 | 98.2 | 95.8 | 97.0 |
| mCherry-mOrange2 | 99.7 | 95.5 | 97.6 |
| Sapphire-mCherry | 99.8 | 91.4 | 95.4 |
| mAmetrine-mCherry | 99.2 | 86.5 | 92.4 |
| LSSmOrange-mCherry | 99.5 | 92.9 | 96.1 |
| EGFP-mCherry | 99.7 | 94.3 | 96.9 |
| EYFP-mCherry | 99.4 | 96.4 | 97.8 |
| mPapaya-mCherry | 99.7 | 97.5 | 98.6 |
| mOrange2-mCherry | 99.5 | 97.6 | 98.5 |
| mCherry-mCherry | 99.0 | 96.6 | 97.8 |
| Mean | 98.7 | 92.8 | 95.6 |
| Median | 99.2 | 94.8 | 96.9 |

To better understand the residual errors, we analyzed the confusion matrix comparing predicted and true barcode identities. Most pairwise misclassifications occurred at rates below 1%, indicating minimal overlap between barcodes. A few specific pairs accounted for higher rates of confusion, including EGFP–Sapphire misidentified as EGFP–EGFP (4.8%) and Sapphire– LSSmOrange misidentified as mAmetrine–mCherry (4.2%) (Fig. 2E). Notably, these errors were asymmetric, with reverse misclassifications occurred at rates below 1%. This low level of classification errors are unlikely to introduce systematic biases in downstream analyses.

### Performance of pooled assays

We next evaluated RainBar classification in pooled settings, where neighboring cell interferences and variable expression levels across constructs could degrade classifier performance. We assembled two representative test pools: one in which all barcodes shared the same nuclear marker (mPapaya–NLS-X) and another with a common cytoplasmic FP (NLS–X-EYFP–NES). When projected onto the supervised UMAP trained on single-barcode cells, pooled cells from both mixtures clustered cleanly with their corresponding references (Fig. 2F,G). Overall classification accuracy remained strong in both pools, with 91.1% and 92.8% for the mPapaya- and EYFP-based sets, respectively, correctly assigned to one of the eight RainBars in the pool, and 8.9% and 7.2% misassigned to other RainBar. These results suggest that the MLP classifier generalizes well across pooled and arrayed contexts. Representative pooled-cell images shown in Fig. 2H.

### Establishing NF-κB live-cell translocation assays

To evaluate RainBar in a biologically relevant setting, one where time-resolved imaging reveals information not accessible through static measurements, we turned to the NF-κB signaling pathway. This pathway is highly dynamic: nuclear translocation of NF-κB subunits can oscillate in some contexts and remain sustained in others^30–32^. Recovery after nuclear entry is mechanistically distinct and tightly regulated while single-cell responses are notoriously heterogeneous. Moreover, NF-κB acts as a hub for signal integration and crosstalk across inflammatory, stress, and developmental inputs (Fig. 3A)^33,33^ These properties make it an ideal testbed for a live-cell imaging platform.

**Figure 3.**
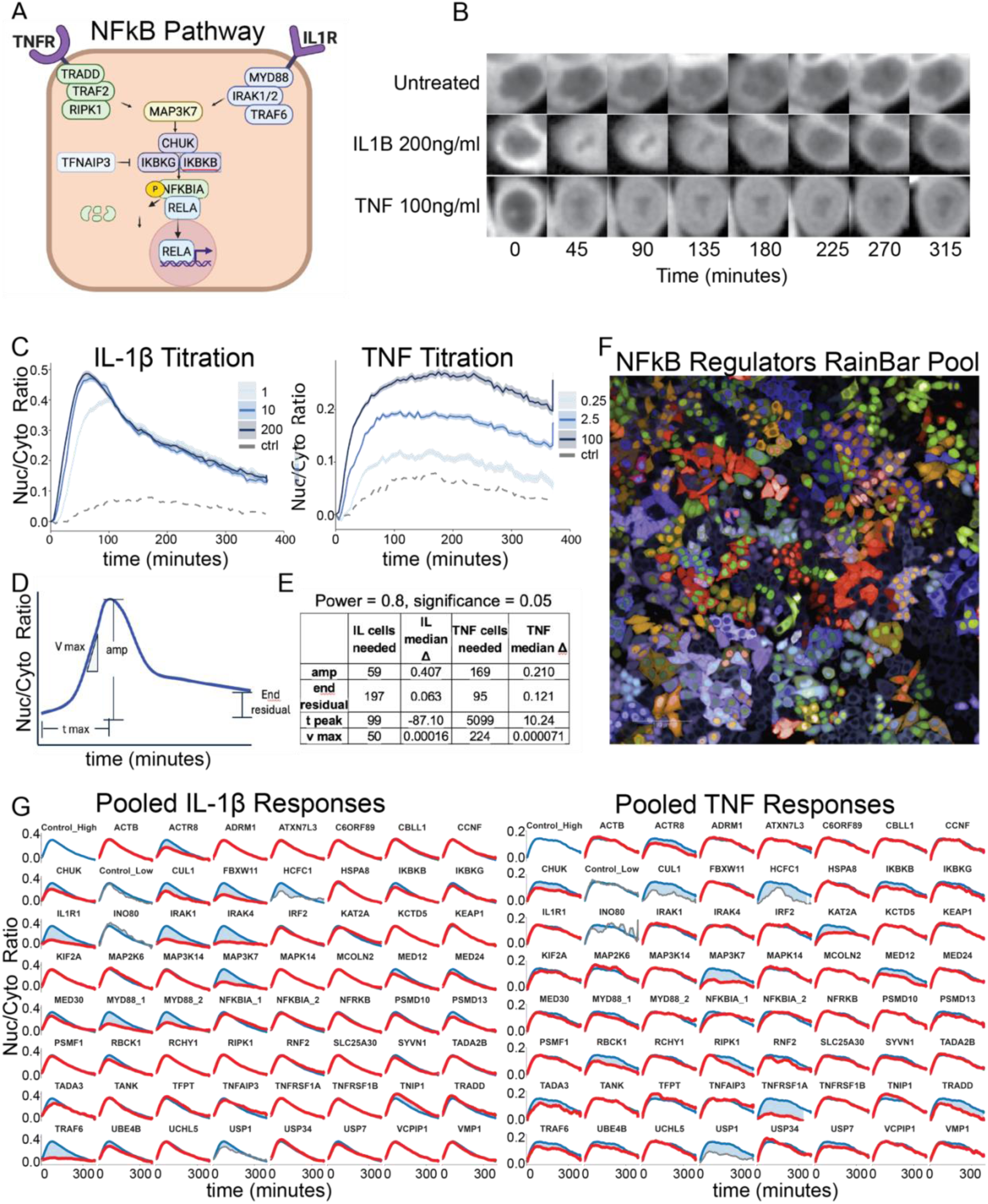
RainBar enables pooled live-cell profiling of NF-κB translocation dynamics. **a,** Schematic of canonical NF-κB signaling pathways downstream of TNFR and IL1R, highlighting both shared and pathway-specific components. **b,** Representative single-cell time-lapse images of HCT116 CRISPRi cells expressing RelA–HaloTag, BFP-NES (segmentation), and barcoded sgRNAs, following stimulation with control, IL-1β, or TNF. Nuclear accumulation of RelA is visible across frames. **c,** Mean nuclear-to-cytoplasmic (N/C) RelA ratio over time for increasing concentrations of IL-1β or TNF. Untreated cells show mild baseline translocation due to phototoxicity from violet excitation. **d,** Schematic of the four kinetic features extracted from each single-cell trace: amplitude (max N/C), time-to-peak (t_peak_), maximum slope (v_max_), and residual nuclear signal at 6 hours. **e,** Power analysis estimating the number of cells required per construct to detect a statistically significant difference (α = 0.05, power = 0.8) for each kinetic parameter under IL-1β or TNF stimulation. **f,** Representative field of view from pooled NF-κB assay. **g,** Mean N/C RelA trajectories for each perturbation (red) under IL-1β (left) and TNF (right), overlaid on the non-targeting control (blue). Perturbations with insufficient cell counts for at least one metric (per power analysis in e) are shown in gray. Shading denotes deviation from control.

We engineered a CRISPRi-compatible HCT116 cell line by inserting a Zim3–dCas9–BFP cassette into the AAVS1 locus. While the BFP signal was detectable by flow cytometry, it was dim enough to avoid interference in imaging-based assays (Fig. 1C)^34,35^. CRISPRi activity was confirmed via CD81 knockdown (Fig. S3A). We then introduced two additional components: BFP-NES-Hygromycin for cytoplasmic segmentation and RelA–HaloTag–Zeocin with JF-646 labeling for real-time tracking of NF-κB nuclear localization^20,24^. Nuclei were tracked over time using the Trackpy particle tracking library and nuclear and cytoplasmic fluorescence intensities were extracted frame by frame^36^. To minimize edge artifacts, we applied a 2-pixel erosion to both cytoplasmic and nuclear masks.

Upon stimulation with IL-1β or TNF, we observed RelA translocation into the nucleus, as expected (Movie 1; Fig. 3B,C,S3B). We quantified four dynamic metrics: (i) maximum nuclear-to-cytoplasmic ratio (N/C), (ii) maximum slope of nuclear entry (v_max_), (iii) time to peak translocation (_tpeak)_, and (iv) residual nuclear signal after 6 hours (Fig. 3D). Because cells in the same well were imaged up to 5.5 minutes apart, we linearly interpolated timepoints for alignment. To preserve raw signal properties, smoothing was applied for visualization only and not for metric extraction.

Interestingly, even untreated cells exhibited low-level nuclear localization of RelA, likely due to phototoxic stress from repeated violet laser exposure (Fig. 3C). The effect was more pronounced with 425 nm excitation than 405 nm, in line with earlier observations^27^. While this low-grade activation was tolerable for NF-κB, more light-sensitive pathways may benefit from red-shifted segmentation markers or label-free segmentation strategies. Notably, debarcoding requires higher-intensity imaging, which itself induces NF-κB activation; we therefore recommend performing functional imaging assays prior to barcode classification.

Kinetic responses to stimuli differed markedly. IL-1β induced rapid nuclear import followed by recovery toward baseline, while TNF triggered more sustained translocation. We did not observe oscillatory behavior in HCT116 cells, consistent with prior reports in epithelial contexts (Fig. 3C)^30,31,37^

To guide pooled screen design, we conducted power analyses to estimate the number of cells required per perturbation to reliably detect kinetic differences (Fig. 3E). For most metrics, ∼225 cells per barcode were sufficient (equivalent to ∼14,400 cells per 96-well plate) but tpeak under TNF stimulation required closer to 6,000 cells, likely due to flatter, more ambiguous peak profiles.

### Pooled regulator screen

To demonstrate the utility for RainBar in high-content functional screening, we applied it to a pooled assay targeting known and suspected regulators of NF-κB signaling. The perturbation set was adapted from a previously validated arrayed screen^5^ and guides were cloned into RainBar constructs alongside two internal controls: mOrange2– mCherry (at 1× DNA input) and mCherry–mCherry (at 2×). Lentivirus was produced using the PRECISE system and transduced into HCT116–dCas9–BFP–RelA cells at low multiplicity of infection (MOI < 30%) to reduce multi-guide integration events. Conveniently, three fluorophores (EGFP, EYFP, and mPapaya) spanned roughly 3/8 of the library and could be visually monitored under 488 nm excitation, providing a coarse estimate of MOI.

Stimulated cells were imaged under IL-1β and TNF in a single time-lapse experiment across triplicate wells (Fig. 3F, G). Representation varied across constructs (Fig. S4), but most perturbations yielded sufficient number of cells over the power threshold. Several constructs showed marked depletion (>2 Log2Fold relative to controls), including TNFAIP3, TRADD, MED30, and USP1, consistent with known viability effects. ^1,37^ Notably, the underrepresented mOrange2–mCherry control increased in abundance during analysis, confirming that observed coverage is due to low input rather than poor fitness in control vectors (Fig. 4A).

**Figure 4.**
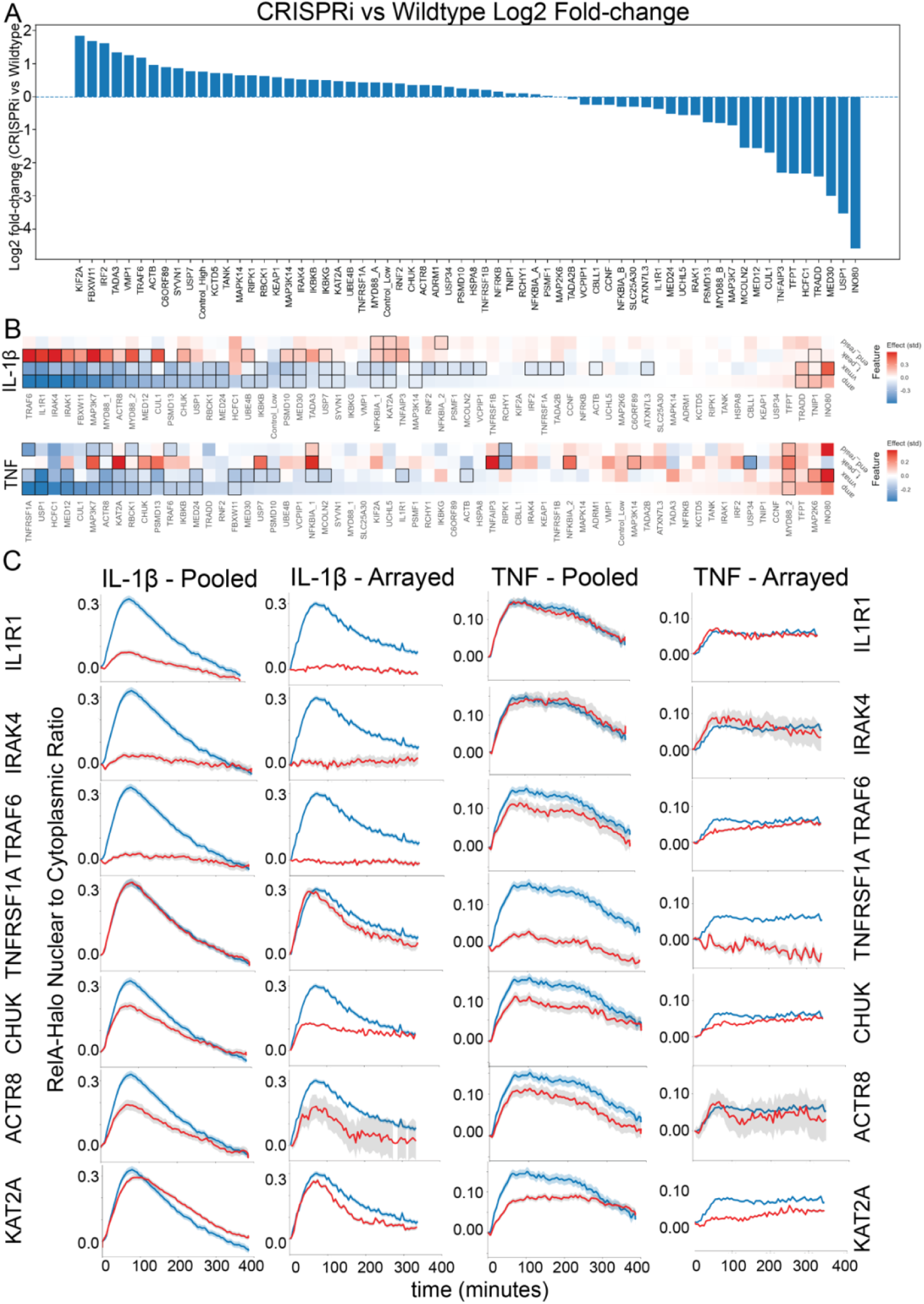
Pooled RainBar screen reveals canonical and candidate regulators of NF-κB dynamics. **a,** Relative abundance of each perturbation in the pooled screen, shown as log₂ fold-change in barcode representation compared to wild-type cells. Essential genes are depleted, while low-concentration control barcodes (e.g., mOrange2–mCherry) increase in relative abundance. **b,** Heatmaps showing effect sizes across four kinetic features, e,g, amplitude, maximum slope (v_max_), time-to-peak (t_max_), and residual nuclear signal, for both IL-1β and TNF stimulation. Perturbations with significant effects (FDR < 0.05) are outlined in black. **c,** Mean RelA N/C trajectories for selected perturbations under IL-1β and TNF stimulation, comparing pooled (RainBar) and arrayed measurements. Shaded regions show 95% confidence intervals. IL-1β regulators (IL1R1, IRAK4, TRAF6), TNF regulators (TNFRSF1A), common regulators ( CHUK) and nuclear factors (ACTR8 and KAT2A) exhibit reproducible, stimulus-specific phenotypes.

Kinetic profiling across debarcoded trajectories revealed diverse effects on NF-κB dynamics. Among the extracted features, v_ₘ_ₐₓ emerged as the most sensitive: it was significantly altered by 31 and 16 perturbations under IL-1β and TNF, respectively, followed by amplitude, t_ₘ_ₐₓ, and residual signal (Fig. 4B). Canonical IL-1β components (IL1R1, IRAK1/4, MYD88, TRAF6) robustly suppressed IL-1β responses, while TNFRSF1A knockdown specifically impaired TNF-induced translocation, as expected^33^. Some regulators such as TRAF6 affected both stimuli responses, pointing to broader roles in inflammatory signaling.

Our live cell RainBar screening also resolved more nuanced phenotypes. Core IKK components (CHUK/IKKα, IKBKB/IKKβ, IKBKG/NEMO) showed partially overlapping functions consistent with their known redundancy. However, subtle kinetic shifts suggested dosage-sensitive effects^33^. Negative feedback regulators, including TNIP1 (ABIN) and TNFAIP3 (A20), modulated both amplitude and termination dynamics, although strong dropout of A20 limited follow-up in the pooled format. Strikingly, chromatin remodelers, especially the Ino80 complex subunit ACTR8, emerged as candidate modulators of NF-κB dynamics, hinting at underappreciated nuclear regulatory layers.

Validation on an independent microscope confirmed the directionality of key hits, despite modest shifts in baseline phototoxicity. IL1R1 and IRAK4 consistently decreased amplitude and v_max_ while delaying t_peak_ under IL-1β. Meanwhile, ACTR8 and KAT2A showed reproducible reductions in amplitude across both stimuli, though the magnitude of effect varied between IL-1β and TNF (Fig. 4C).^5,11^ These results underscore the capacity for RainBar not only to recover known pathway architecture but also to surface new players in dynamic regulation.

## DISCUSSION

RainBar introduces a new strategy for pooled optical screening, enabling multiplexed live-cell imaging of dynamic cellular phenotypes. By addressing long-standing technical barriers, we establish that fluorescent barcodes can be reliably delivered, decoded, and used in complex assays at scale. These include: lentiviral recombination segmentation of both nuclear and cytoplasmic compartments and resolution of spectrally overlapping fluorophores. The current 64-barcode set supports medium-throughput screens and is readily expandable across multi-well formats for genome-wide applications.

To illustrate the utility of RainBar, we profiled NF-κB signaling, a pathway well known for its rich temporal complexity that challenges common endpoint readouts. Our results confirmed core pathway regulators, including IL1R1, MYD88, and TNFRSF1A, while also uncovering more subtle phenotypes. For instance, IKK subunits displayed partial functional redundancy with dose-sensitive kinetic effects, and negative regulators such as TNIP1 and A20 modulated termination dynamics. Chromatin remodeling factors, ACTR8, also emerged as candidate modulators, pointing toward potentially unexplored regulatory crosstalk between nuclear architecture and inflammatory signaling.

Several advantages distinguish RainBar from existing pooled screening approaches. Chief among them is the ability to perform assays in living cells, capturing real-time signaling behaviors that static methods overlook. The platform is compatible with orthogonal readouts, such as HaloTag reporters and avoids the destructiveness of sequencing-based endpoints. That said, some limitations persist: violet laser excitation introduces phototoxic stress over long time courses, essential genes may be underrepresented due to viability effects, pooled dropout, and scaling beyond 64 perturbations requires multi-plate designs or broader localization strategies. These limitations are not unique to RainBar but rather reflect broader tradeoffs inherent in optical pooled screening.

Looking forward, RainBar opens the door to diverse applications, from dissecting nuclear transport and transcriptional burst kinetics, to screening combinatorial perturbations or real-time feedback mechanisms in signaling networks. By enabling systematic, multiplexed observation of dynamic biology at single-cell resolution, it offers a flexible platform for uncovering how molecular circuits operate– not just on average, but in motion, in context, and in real time.

## METHODS

### Cell lines and culture

HEK293T (ATCC CRL-3216) and PRECISE producer cells were maintained in Dulbecco’s Modified Eagle Medium (DMEM, Gibco 11965-092) supplemented with 10% fetal bovine serum (FBS, Gibco 26140-079) and 1% penicillin–streptomycin (Gibco 15140-122). HCT116 cells were short tandem repeat (STR)-certified and obtained from the UCSF Cell and Engineering Core, and maintained in McCoy’s 5A medium (Gibco 16600-082) supplemented with 10% FBS and 1% penicillin–streptomycin. All cell lines were grown at 37 °C in a humidified atmosphere with 5% CO₂.

### Lentivirus

Lentiviral particles were produced in HEK293T or PRECISE cells using Lipofectamine 3000 (Thermo Fisher L3000015) and third-generation helper plasmids psPAX2 (Addgene 12260) and pMD2.G (Addgene 12259). Viral supernatants were used directly for transduction without concentration or polybrene to maintain low multiplicity of infection (MOI). Target cells were transduced with RainBar constructs and imaged 10–15 days post-transduction to ensure stable expression.

### NF-κB assays

For live-cell assays, HCT116 cells were plated in 96-well format and stimulated with recombinant human IL-1β (InvivoGen, cat. #rhu-il1b) or TNF (InvivoGen, cat. #rhu-tnfa) at the concentrations specified in the main text and figure legends. Imaging began within 2 min of cytokine addition, and time-lapse acquisition proceeded for 65 frames. Nuclear DNA was counterstained with Hoechst 33342 (Thermo Fisher H3570) and/or SPY700-DNA (Spirochrome SC501) following manufacturers’ recommendations.

### Imaging

Time-lapse imaging was performed using Opera Phenix high-content screening systems. NF-κB assays and debarcoding experiments requiring additional subfilters were conducted at the UC Berkeley High-Throughput Screening Facility on a 5-laser Opera Phenix (405, 425, 488, 561, 640 nm). All other experiments were conducted at UCSF on a 4-laser Opera Phenix (405, 488, 561, 640 nm). Laser intensities and exposure times are listed in Supplementary Table 1. Vector construction and codon optimization

### Cloning

Final RainBar vectors were ordered at plasmids from Twist biosicences. sgRNA and barcode elements, were synthesized by IDT and cloned as previously described^38^. Fluorescent protein coding sequences were codon-optimized manually to balance expression efficiency and minimize nucleotide-level homology between nuclear and cytoplasmic variants, while retaining amino acid identity.

### Barcode classifier (MLP)

Training. Nuclei and cytoplasm were segmented with Cellpose (v2.2) using HCT116-refined models (estimated diameter 30 px). Cytoplasmic masks were generated from BFP-NES, and nuclear masks were obtained by inverting the same images: the inverted contrast renders nuclei lighter than background, allowing Cellpose to delineate nuclear boundaries accurately. Single-construct wells were segmented and quantified, yielding per-cell mean intensities in 14 channels for nucleus and cytoplasm. Features were built as all pairwise within-compartment ratios, concatenated across nucleus and cytoplasm. Cells with cytoplasm area <10 px or circularity ≥0.8 were excluded.

Location specific channel ratios were calculated, labels encoded, and split 70/30 (seed 123). We trained an MLP (4 layers, 64 nodes/layer) with Adam and cross-entropy for 10 epochs (batch 32). Reported metrics included held-out accuracy, recall, and F1 score. Fitted scaler/encoder/model were saved for deployment.

For visualization and to generate reusable backgrounds for pooled overlays, the penultimate layer of the trained MLP (“likelihood” all 64 barcodes X number of cells) was extracted, this 64 X n matrix was dimensionality reduced with PCA to give a PCA embedding. This PCA embedding was further dimensionality reduced with UMAP to yield the 2 X 2 embedding. The MLP embedding, PCA embedding, and the UMAP embedding were all fitted with the training data and then saved. The UMAP embedding was fit on a balanced subset of up to 500 cells per class, filtered to include only high-quality cells (predicted confidence > 0.8). The experimental data was then run though the fixed/saved MLP>PCA>UMAP.

### Processing Pooled Assay Images application to pooled data

Pooled wells were processed identically to generate ratio features, standardized with the training scaler, and classified by the saved MLP to assign RainBar IDs and confidences per cell. Pooled-well QC excluded cells with cytoplasm area <100 px. Images were organized into time-ordered stacks per position/channel, and microscope metadata were parsed to obtain frame numbers and absolute time. Composite stacks overlaying assay fluorescence with nuclear/cytoplasmic boundaries supported visual QC.

Each nucleus was paired to its enclosing cytoplasmic mask (most-overlap rule). A “ring” region was defined as cytoplasm surrounding that nucleus with the nuclear core excluded; ring pixels inherited the nucleus ID to preserve one-to-one matching. Additionally, we eroded 2 pixel from the boundaries to avoid signal from adjacent compartment. For each nucleus–ring pair we computed mean and SD intensities for both compartments and recorded the nuclear centroid. Cells 50–900 px were retained. Cells on the border of the field of view and low-signal cells with (mean nucleus + mean cytoplasm) <700 a.u. were excluded, typically retaining ∼70–80% of objects.

Nuclear centroids were linked across 65 frames with TrackPy (BTree search; search range 10 px; minimum track length 10 frames)^36^. Global drift was corrected by phase cross-correlation when needed. Per frame, N/C was computed with error propagation from compartmental SDs. Missing frames were linearly interpolated. Per-cell records were annotated with well, treatment, and replicate from plate maps. Barcode masks were registered to assay masks via centroid-based deformable matching with a 60-px displacement cap, using Coherent Point Drift (CPD) deformable point-set registration on nuclei centroids, then nearest-neighbor matching with a distance cutoff[37]. Each tracked trace was assigned a RainBar ID and mapped to the corresponding gene.

### R-based analysis of NF-κB dynamics

Single-cell N/C time series and metadata (treatment, plate, well, barcode→gene) were loaded per experiment. Time was converted to minutes and traces smoothed with a centered rolling median (k=5) with mean-fill only where the median produced edge NAs. A per-cell baseline (mean N/C over 0–5 min) was subtracted to obtain nc_bl, used unless noted. From nc_bl we derived: amplitude (max), time-to-peak (t_peak_), maximum slope (v_max_ = ΔN/C/Δt), and end residual (final value).

Each nc_bl trace with ≥8 points was fit to a difference-of-exponentials (“impulse”) using Levenberg–Marquardt. Parameters were baseline BBB, amplitude AAA, rise/decay time constants, and onset t₀ (positive parameters fit in log-space). We recorded RMSE, predicted tₚₑₐₖ, and peak height; BBB served as the base feature.

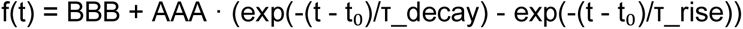

For each treatment × feature (amp, t_peak_, v_max_, end_resid, base), we compared every gene to the non-targeting control using a plate/well–stratified permutation test (B=5000; mean difference, or median for t_peak_). P-values were BH-adjusted within treatment × feature. Where specified, stratified bootstrap CIs (B=2000) were computed for effect sizes.

Per-cell smoothed traces were linearly interpolated to a 0.1-min grid and aggregated to mean ±95% CI per gene/treatment; bins with <10 cells were suppressed. Non-targeting control was overlaid (dashed) in each facet, with y-limits chosen from the CI envelope and padded by 5%.

## DATA AVAILABILITY STATEMENT

### Data availability

The datasets generated during this study will be deposited in AWS and made publicly available upon acceptance.

### Code availability

All code used for image segmentation, barcode decoding, and NF-κB kinetic will be available on GitHub

## ACKNOWLEDGEMENTS

We thank members of the McManus and Huang laboratories for insightful discussions, technical support, and feedback throughout the development of this work. This study was supported, in part, by National Institutes of Health grants U01CA272546 and U24DK116214 to M.T.M., as well as a Laboratory of Genomics Research Innovation Award to M.T.M. and B.H.. M.T.M. and B.H. are Chan Zuckerberg Biohub San Francisco Investigators. This work was assisted by the UCSF Center for Advanced Technologies and is supported by the Diabetes Center at UCSF

## AUTHOR CONTRIBUTIONS

R.M., B.H., and M.T.M conceived the project. D.F., L.S., R.M., and J.F. developed computational pipelines. R.M., D.F., L.S. C.T., N.T., M.G.K., S.O., B.S. and designed and performed experiments and analyzed data. R.M., B.H., and M.T.M supervised experimental design and analysis and advised on assay development. R.M., B.H., and M.T.M wrote/edited the manuscript. All authors reviewed and approved the final manuscript.

---

**Figure S1.**
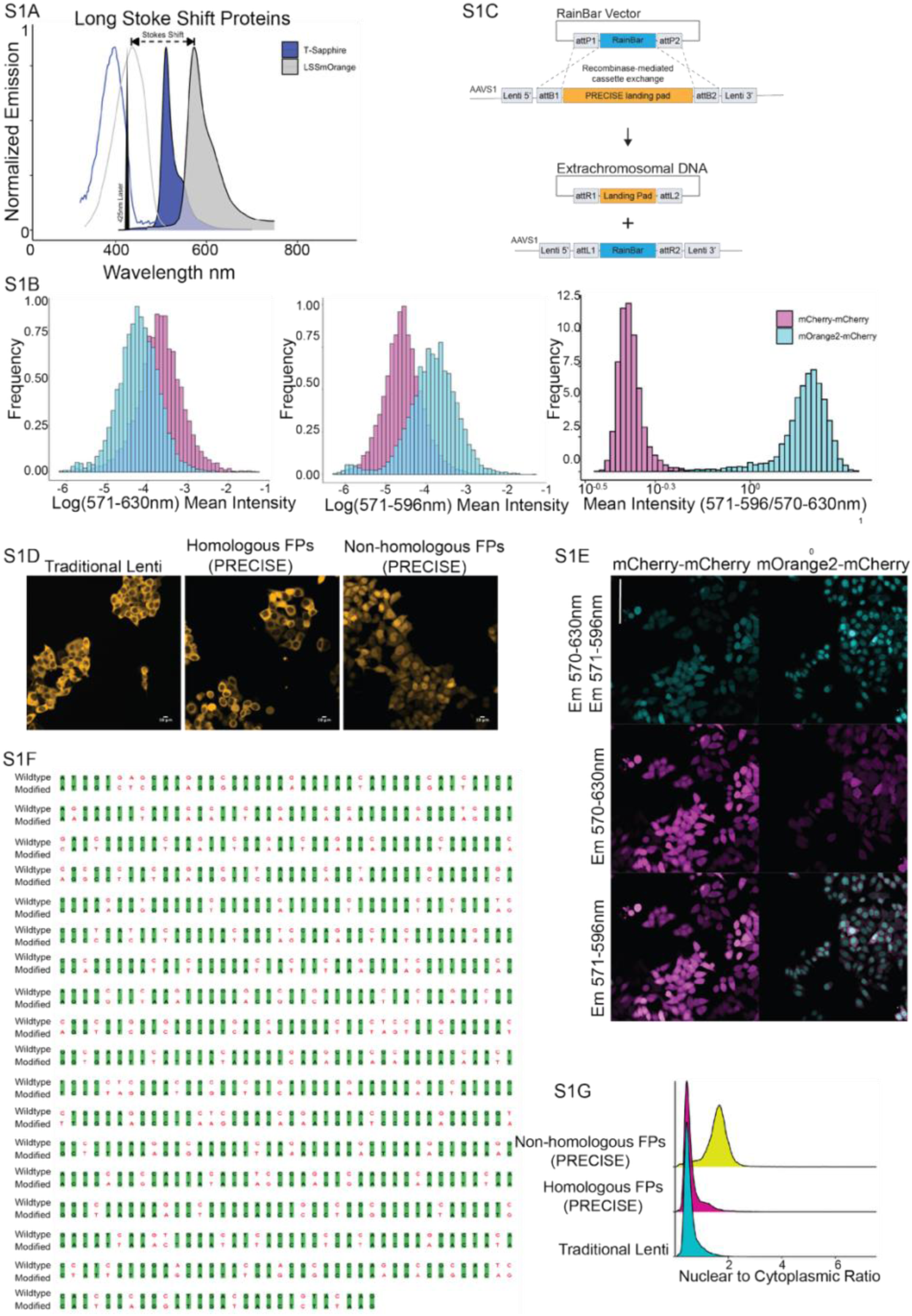
Spectral unmixing and mitigation of lentiviral recombination through codon diversification and single-template production. **a,** Emission profiles of long Stokes shift fluorescent proteins (FPs) Sapphire (EGFP like emission) and LSSmOrange (mOrange2 like emission), both excited at 425 nm. **b,** Fluorescence intensity distributions for dual mOrange2–mCherry and mCherry– mCherry constructs imaged across two emission channels. Histograms (left and middle) show per-cell intensities; right, histogram of log₂ intensity ratios enables separation of spectrally overlapping fluorophores. **c,** Schematic of the PRECISE lentiviral system: a recombinase-addressable landing pad integrated at the AAVS1 locus permits stable integration of a single RainBar cassette per packaging cell. Single-template virus reduces recombination relative to standard pooled transfection. **d,** Representative images of HCT116 cells transduced with mOrange2–mOrange2 constructs generated using standard pooled lentivirus (left), PRECISE with homologous FPs (middle), and PRECISE with codon-diversified NLS-FP (right). **e,** Representative images of condon diversified mCherry-mCherry and mOrange2-mChery demonstrating ability to separately similar colors and lack of recombination artifact **f,** Pairwise alignment of canonical and codon-diversified mOrange2 sequences highlighting reduced nucleotide identity across the coding region. **g**, Nuclear-to-cytoplasmic (N/C) intensity ratios for mOrange2 in cells from panel d. Codon diversification reduces recombination-induced loss of nuclear signal. N/C >1 indicates correct localization.

**Figure S2.**
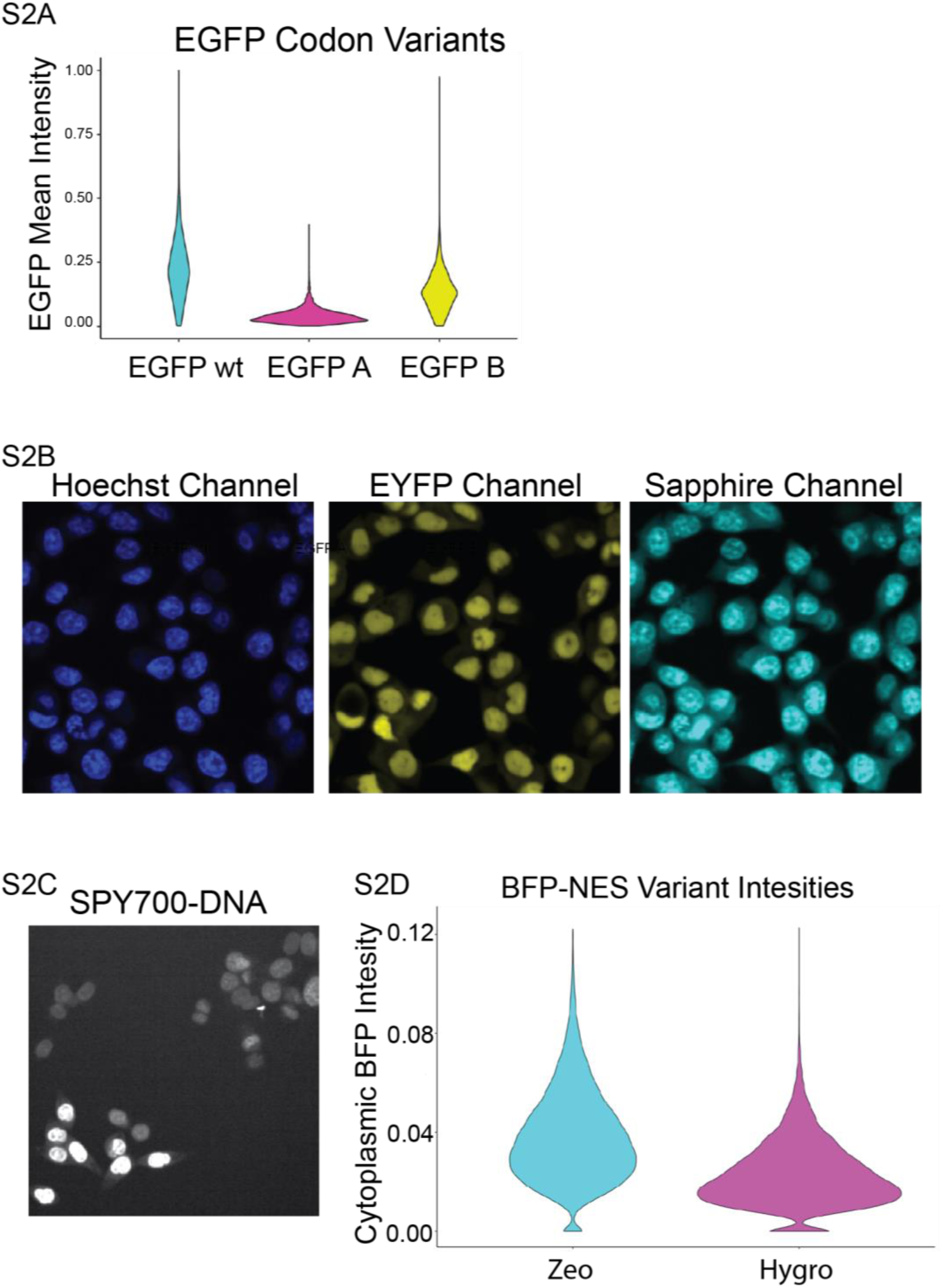
Optimization of fluorescent protein expression and nuclear staining strategies. **a,** Signal intensity of EGFP variants: violin plots show nuclear intensity distributions for canonical EGFP (EGFP wt) and two codon-diversified constructs (EGFP A, EGFP B); the EGFPB variant achieves restored expression compatible with imaging applications. **b,** Spectral interference from Hoechst 33342 in the Sapphire channel. Cells expressing an EYFP–Sapphire RainBar construct show cytoplasmic signal overlap in both the EYFP and Sapphire channels, complicating segmentation and decoding. **c,** SPY700-DNA stain yields uneven nuclear labeling in HCT116 cells, limiting its utility for segmentation. **d,** Signal intensity of BFP–NES constructs encoding either Zeocin or Hygromycin resistance genes. Zeocin yields higher BFP intensity, while the dimmer Hygro variant reduces bleed-through into neighboring channels.

**Figure S3.**
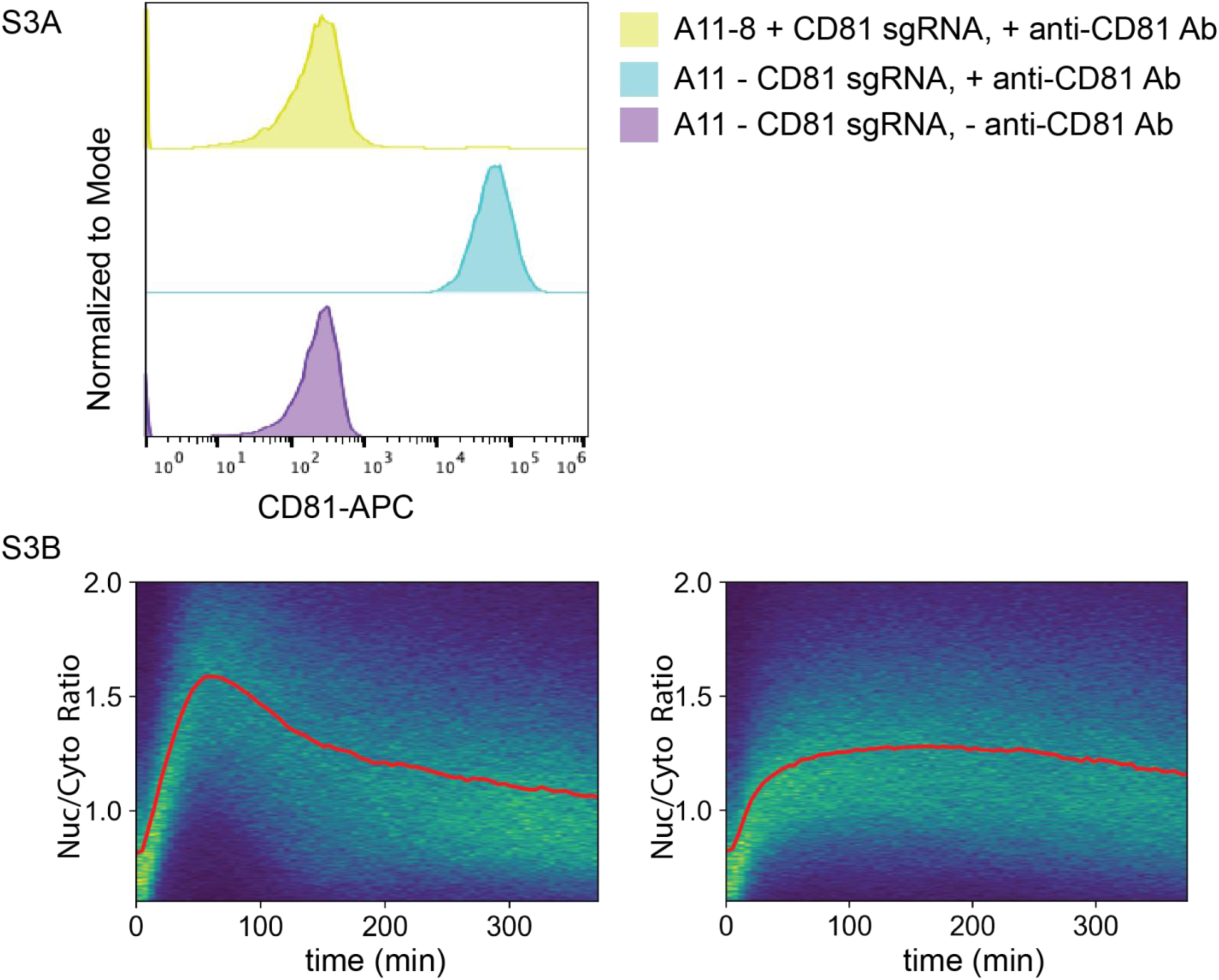
Assay validation. **a,** Validation of Zim3–dCas9–BFP CRISPRi system. Clone A11-8, engineered to express BFP–NES and RELA–HaloTag, shows efficient knockdown of CD81 relative to the A11 unstatined parental cell. **b,** Density plot showing single-cell heterogeneity in RelA–HaloTag translocation following stimulation. Raw nuclear-to-cytoplasmic (N/C) traces shown as density heatmaps for IL-1β and TNF; red lines indicate the mean trace.

**Figure S4.**
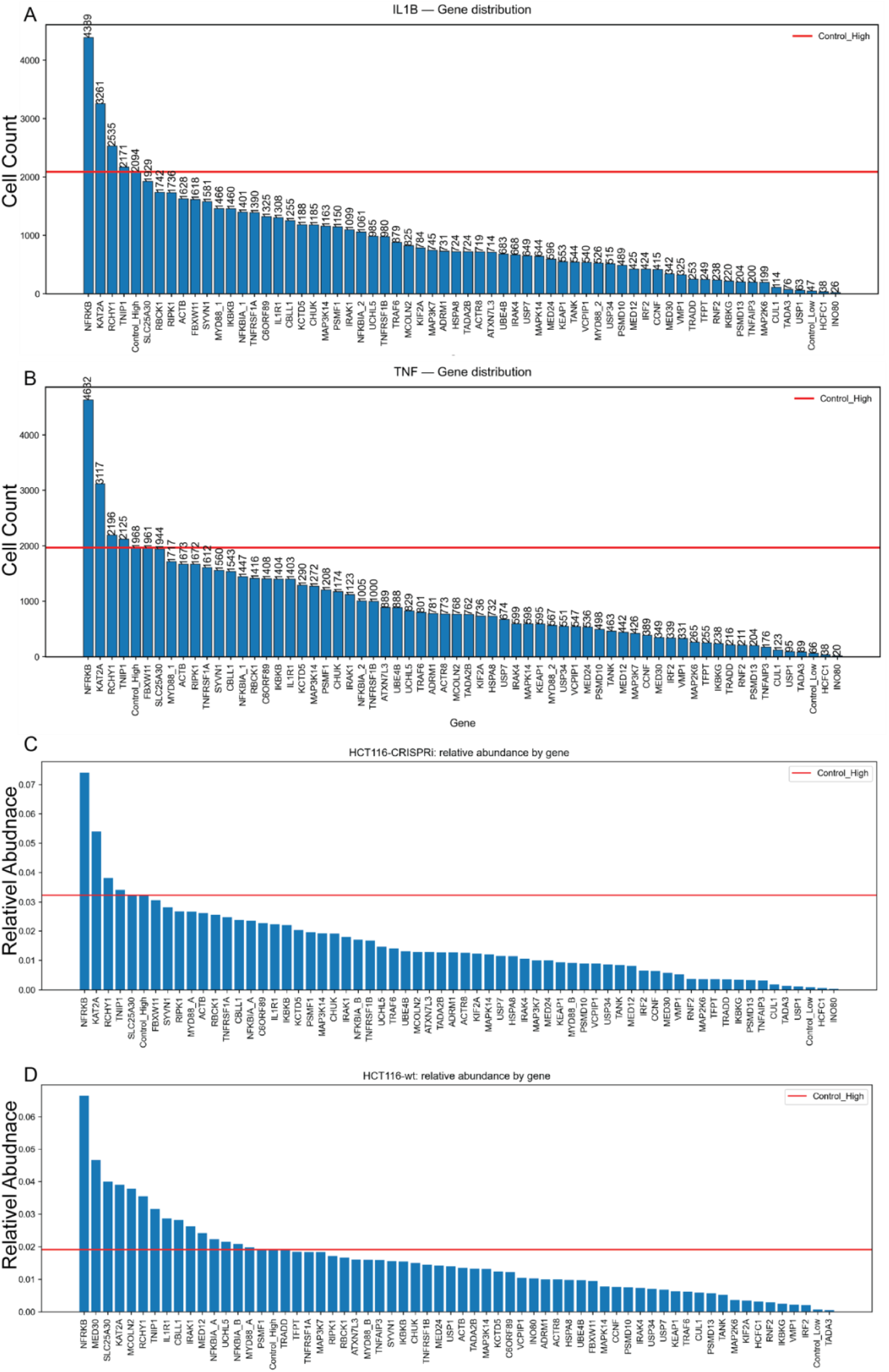
Construct representation in pooled screens. Construct representation in pooled experiments. **a** and **b,** gene-level cell counts for each perturbation under IL-1β and TNF, respectively. **c** and **d,** relative barcode abundance in HCT116-CRISPRi and wildtype HCT116 cells, respectively, normalized across the pool. Reductions in representation reflect low transduction or cell fitness effects.

